# Bestrophin1-mediated GABA release activates large chloride currents to generate epileptiform events in the entorhinal cortex

**DOI:** 10.1101/2025.04.15.648911

**Authors:** Paolo Scalmani, Laura Uva, Maria Cristina Regondi, Claudia Miele, Valentina Grazioso, Marco de Curtis

## Abstract

The mechanisms that lead to the onset of a focal seizure are still not understood. We used the well-established 4-aminopyridine (4AP) ictogenesis model to analyze epileptiform discharges in the entorhinal cortex of rodents maintained *in vitro*. Simultaneous field potential and patch-clamp recordings demonstrated that 100 µM 4AP elicited periodic and large chloride currents in both principal neurons and GABAergic interneurons that steadily matched with population spikes. These *population spike-associated chloride currents* (*PSACCs*) survived glutamate and glycine receptor blockade and were abolished by GABA_A_ antagonists and by blocking synaptic neurotransmitters release. Antagonist of astrocyte bestrophin-1 channels inhibited PSACCs and prevented the occurrence of seizure-like events in both entorhinal cortex mouse slices and in the isolated guinea pig brain.

We propose that bestrophin-1-induced GABA release likely triggered by astrocytes promotes in all neurons subtypes a large chloride current that is responsible for interictal spikes and establishes the conditions for the generation of seizure-like events.

## Introduction

Neuronal discharges within cortical networks are responsible for the generation of synchronous activities that represent the foundation of physiological brain rhythms. Pathological interactions within an altered epileptogenic network can trigger epileptiform interictal events and focal seizures. Despite the vast literature on the topic, it is not yet established what are the mechanistic determinants that promote the generation of focal seizures. To approach this question, cellular and network neurophysiological patterns observed during epileptiform events that mimic human focal epileptiform discharges have been analyzed on experimental preparations. Neuronal activity patterns recorded during focal seizures in different *in vitro* and *in vivo* rodent models and in humans were recently analyzed, compared and discussed^1^; this report showed that the participation of both principal excitatory neurons and γ-aminobutyric acid-ergic (GABAergic) interneurons can be observed (seldom simultaneously) in diverse models of focal ictogenesis and in single unit recordings performed during intracerebral presurgical monitoring in human focal epilepsies^2,3^.

One of the best established and characterized *in vitro* method utilized to experimentally induce focal seizure-like events (SLEs) is the perfusion of cortical tissue slices with low Mg^2+^ solution added with the potassium channel blocker, 4-aminopyridine (4AP)^4,5^; the slowing of action potential repolarization induced by 4AP prolongs presynaptic terminal depolarization and increases the presynaptic release of neurotransmitter^6–8^. When this model is applied to *in vitro* preparations, entorhinal cortex (EC) and hippocampal SLEs are reliably preceded by heading population spikes that correlate with the paradoxical activation of GABAergic interneurons^9–16^ paired with a rise in the extracellular potassium^17,18^. We demonstrated that ictal SLEs in the EC are abolished by blocking either the glutamatergic or the GABAergic synaptic transmission with kynurenic acid and picrotoxin, respectively^19^; moreover, 4AP-induced interictal epileptiform spikes resisted glutamatergic receptor antagonism and were abolished by GABA_A_ receptor blockers^20^. More recently, we studied the specific entrainment of different GABAergic interneuron subtypes of the mouse EC at the onset of 4AP-induced SLEs^19^. Using patch clamp recordings coupled with single cell digital PCR analysis, we demonstrated that GABAergic cells expressing either cholecystokinin, parvalbumin or somatostatin were recruited at the very beginning of a SLE, either before or together with principal pyramidal cells. We also observed that all neuron subtypes (both principal cells and interneurons) were active during the preictal heading spikes that precede SLEs. The above experiments demonstrated that GABAergic networks are involved in 4AP-induced EC ictogenesis, but do not clarify how these neurons are synchronously recruited.

To further analyze the possible generators of synchronous ictal and preictal epileptiform events in the 4AP model, we performed a detailed pharmacological investigation of synaptic and extrasynaptic channels and of transporters in mouse EC *in vitro* slices bathed in 4AP. We showed for the first time that a large chloride conductance is generated in both pyramidal neurons (PNs) and GABAergic interneurons (INs) in steady coincidence with the onset of epileptiform population spikes; we also demonstrated that the blockade of GABA release mediated by bestrophin-1 (BEST-1) channels abolished this spike-associated chloride currents and prevented the generation of both population spikes and SLEs generated by 4AP in both EC mouse slices and in the EC of the *in vitro* isolated guinea pig brain.

## Results

Experiments were performed in 120 EC slices obtained from xx C57BL/6J mice that specifically express GFP in GABAergic interneurons (see Methods) and in 5 *in vitro* isolated guinea pig brains. Intracellular patch-clamp recordings were performed from pyramidal neurons (PNs; n=90 selected for the study) and GABAergic interneurons (INs; n=80) simultaneously recorded with extracellular local field potentials (lfp). In 5 experiments, simultaneous double patch-clamp recordings from PN and IN pairs were performed; Figure 1a illustrates the typical activity recorded from an IN recorded in current-clamp setting (upper trace) and from a PN recoded in voltage clamp (middle trace, holding potential at -70 mV; 139 mM intracellular chloride concentration) and the simultaneous lfp (lower trace) recorded at < 1 mm from the recorded neurons, before and during the application of 100 µM 4AP. A reproducible set of events was reliably observed in all recorded PNs and INs^19^: at the onset of 4AP perfusion the background spontaneous activity became larger in amplitude (dash-underscored traces in Fig. 1a) and gradually organized to generate population spikes (asterisks Fig. 1a) that evolved into a SLE headed by a large population spike (open arrow on the right in Fig. 1a). To analyse the mechanisms of pre-ictal events, we focused our attention on the intracellular correlates of 4AP-induced population spikes, that recurred with a 0,075 ± 0,003 Hz periodicity (n=125) when glutamate transmission was blocked with 3 mM kynurenic acid (KYN; asterisks in the right panel in Fig. 1b, and Fig. 2a-c) to abolish SLEs^18,19^.

**Figure 1.**
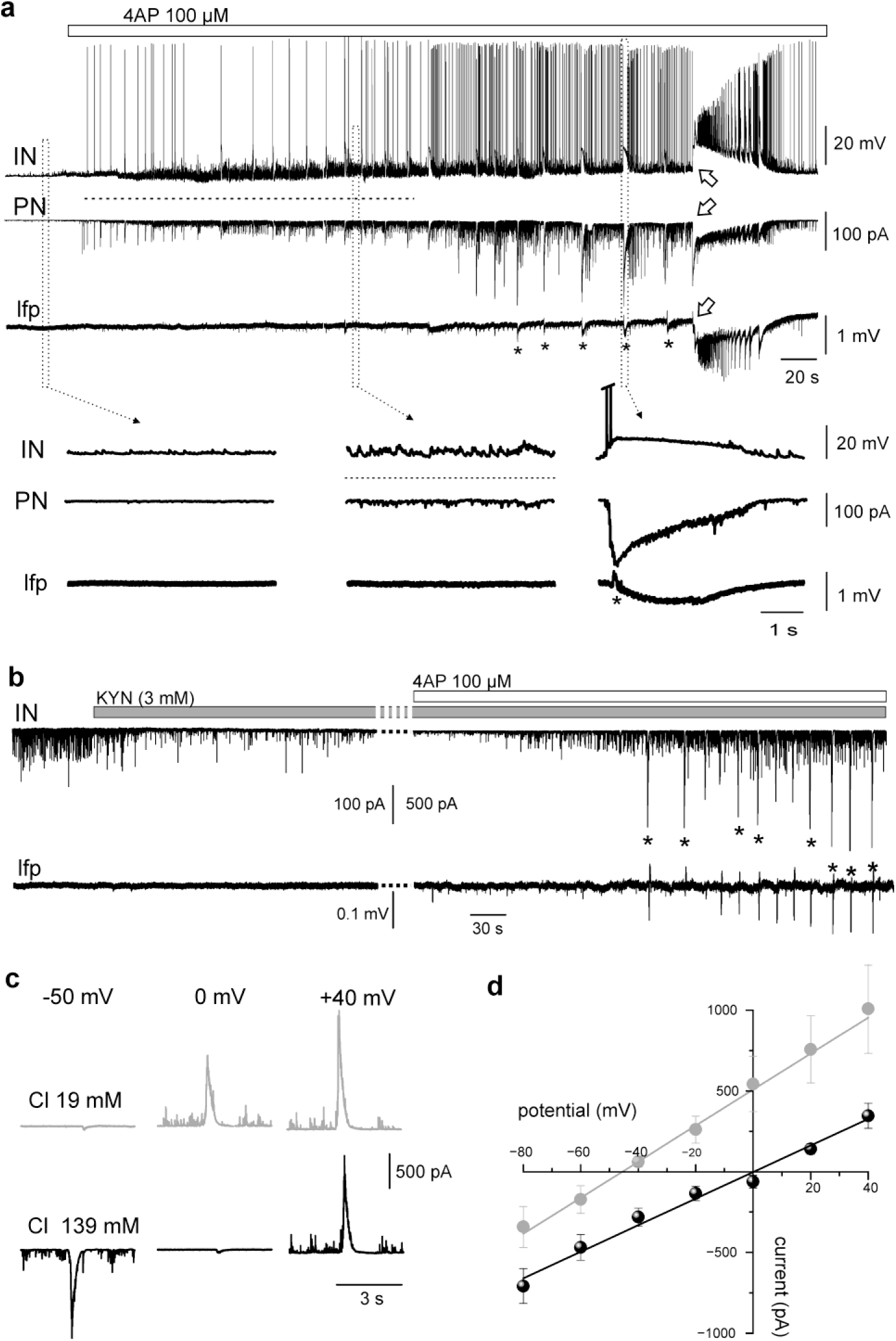
Epileptiform activity in EC mouse slices exposed to 4-aminopyridine (4AP). **a.** Simultaneous extracellular local field potential recordings (lfp, lower trace) and double patch recording from an EC interneuron (IN; upper trace) and pyramidal neuron (PN; middle trace) during slice perfusion with 100 µM 4AP (white bar on top). The signals outlined in the boxes are illustrated with expanded time scale in the lower part of the figure. Lfp population spikes are marked by asterisks; the arrow points to the onset of the seizure-like epileptiform discharge. Note the enhanced background activity induced by 4AP, highlighted by the dash-underscored horizontal line. **b.** Voltage clamp recording from an IN (upper trace) and simultaneous extracellular lfp (lower trace) before and during slice application of 100 µM 4AP (white bar on top), co-perfused with 3 mM kynurenic acid (KYN; gray bar). KYN abolished glutamatergic background activity. The asterisks indicate the population spikes associated with large amplitude inward currents. Note the difference in the intracellular trace calibration between the left and right panel. **c**. Voltage clamp recordings of the reversal potential of the large amplitude inward conductance induced by 4AP during KYN-4AP co-perfusion: recordings were performed either with standard chloride concentration in the recording patch pipette (19 mM; upper grey traces) or with high chloride concentration (139 mM; lower black traces). From left to right, the voltage of the neuron membrane was clamped at -50 mV, -0 mV and +40 mV. **d.** Average current/voltage plots of the slow current obtained in patch clamp experiments performed with standard (19 mM chloride, grey line and symbols, n=9) and high chloride concentration (139 mM; black line and symbols, n=13) in the intracellular pipette solution.

**Figure 2.**
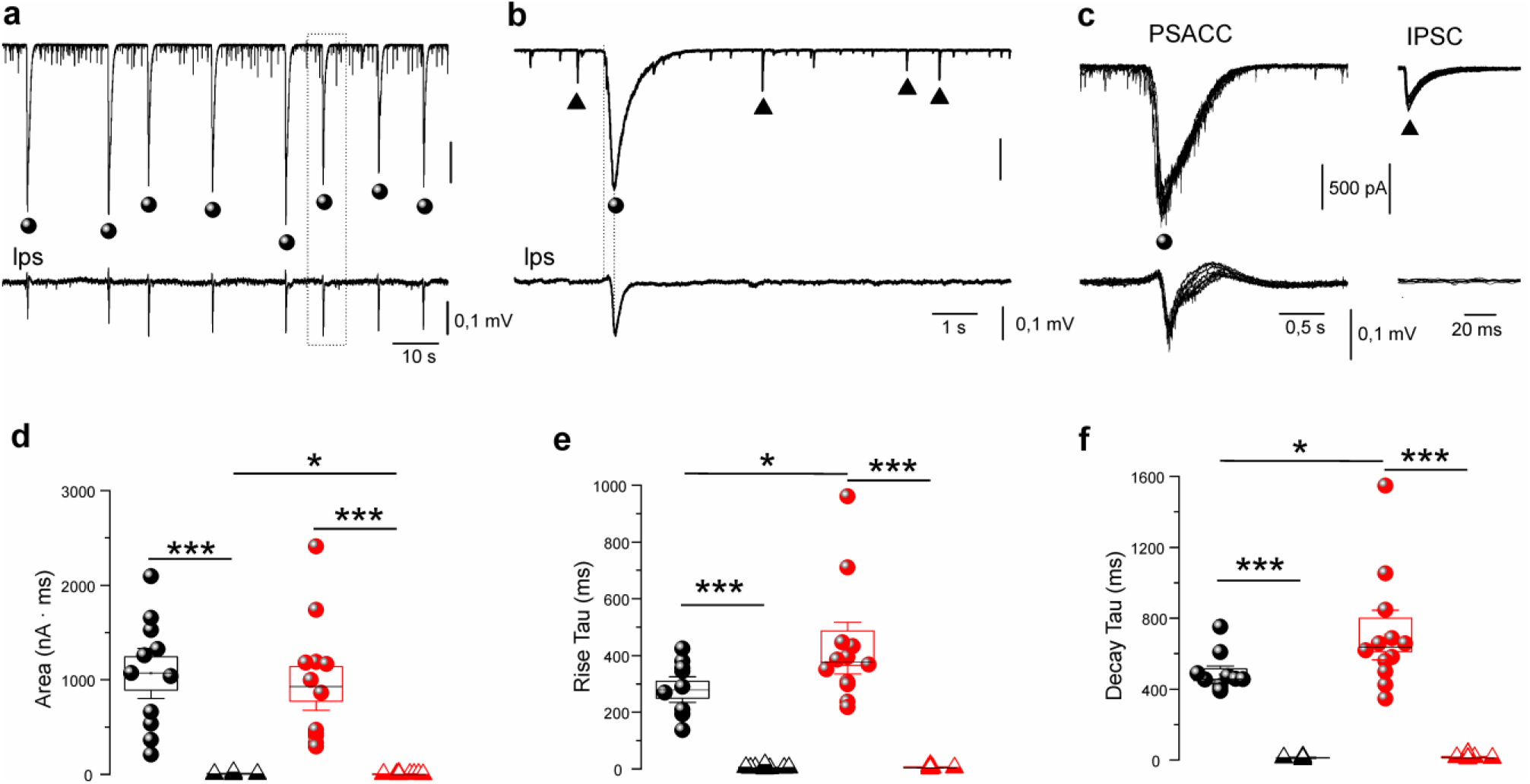
Slow and fast chloride current kinetics. **a**. Slow and large chloride conductances (spheres) and fast chloride currents (triangles) recorded from an interneuron during slice perfusion with 3 mM KYN and 100 µM 4AP. **b.** Time magnification of the slow (spheres) and fast chloride currents (triangles) outlined by the dotted box in a. **c.** Superimposed slow chloride currents (top left traces) and the correlated lfp population spikes (bottom left traces) and fast Cl^-^ conductances (IPSCs; top right traces) obtained during intracellular voltage clamp recording at -70 mV. IPSCs are not correlated with population spikes (bottom right traces). **d-f.** Features of population spike-associated chloride currents (PSACCs; spheres) and fast IPSCs (triangles) recorded from 13 PNs (black symbols) and 11 INs (red symbols). From left to right, quantification of PSACCs and IPSCs area, rise *τau* and decay *τau* for PNs (black) and Ins (red). T-test and Wilcoxon Mann Whitney test were used. Statistical significance: * p<0.05, **p<0.01, *** p<0.005.

### Population spike-associated Chloride Current

As illustrated in representative experiment of Figure 1b, the background spontaneous synaptic activity was reduced by blocking glutamate receptors with either 3 mM KYN (n=10) or a mix of the NMDA (100 µM aminophosphonovaleric acid) and non-NMDA (50 µM 6-cyano-7-nitroquinoxaline-2,3-dione) receptors antagonists (n=10; data not shown). Under these conditions, two types of non-glutamatergic currents with different amplitude and kinetic were consistently observed in both INs and PNs (n=170); the largest current was regularly coupled with a lfp population spikes (spheres in Fig. 2a-c), suggesting that it is simultaneously generated in a large number of cells. Voltage clamp experiments with standard intracellular solution (19 mM chloride concentration in the recording pipette) showed that the slow current had the same reversal potential around -50 mV (n=9) indicative of a chloride (Cl^-^) membrane conductance (Fig. 1c,d). When patch clamp recordings were performed with high Cl^-^ concentration (139 mM) in the recording pipette, the large spike-associated conductance reversal shifted toward more positive potentials (0 mV, n=13; Fig. 1c,d) confirming that this current is carried by Cl^-^ ions. In the following Figures intracellular Cl^-^ concentration of the patched INs and PNs was 139 mM Cl^-^ to enhance PSACC size.

Figure 2 illustrates the main features of the two Cl^-^ conductances recorded in 10 PNs (black symbols in Fig. 2d-f) and 11 INs (red symbols in Fig. 2d-f). Compared to the small-amplitude Cl^-^ currents (triangles in Fig. 2b-f), the ***population spike-associated chloride current*** (***PSACC***; spheres in Fig. 2a-f) was larger and longer over time, presenting slower rise and decay times, as highlighted in Figure 2b-f and in Table 1. Separate analysis of Cl^-^ currents in PNs and INs showed that PSACCs were significantly slower in INs (Fig. 2e and f); the overall current charge (area under the trace) was not significantly different in PNs and INs (Fig. 2d). The small amplitude Cl^-^ currents area was slightly different in the two groups of neurons (compare black *vs* red triangles in Fig. 2d). Table 1 summarizes the main current parameters analysed, that include area, rise *τau*, decay *τau*, time to peak, maximum rise slope and half-width.

**Table 1.** Main features of the two Cl-conductances (PSACCs and IPSCs) recorded in PNs and INs. Values outlined with **bold** character are statistically significant for the indicated paired category, according to the detailed *p* values. MRS= maximal rise slope.

|  |  | Area (nA*ms) | Rise *tau* (ms) | Decay *tau* (ms) | Time to peak (ms) | MRS (pA/ms) | Half-width (ms) |
| --- | --- | --- | --- | --- | --- | --- | --- |
| Cl<sup>-</sup> conductances | PSACCs PN | **1067,06 ± 175,53**<br>*p*<<0.001 vs IPSACs PN | **279,82 ± 30,06**<br>*p*<<0.001 vs IPSACs PN<br>*p*=0,032 vs PSACCs IN | **480,94 ± 32,78**<br>*p*<<0.001 vs IPSACs PN<br>*p*=0,021 vs PSACCs IN | **242,41 ± 12,53**<br>*p*<<0.001 vs IPSACs IN<br>*p*=0,013 vs PSACCs IN | **-40,19 ± 6,28**<br>*p*<<0.001 vs IPSACs PN<br>*p*=0,001 vs PSACCs IN | **514,75 ± 19,44**<br>*p*<<0.001 vs IPSACs PN<br>*p*=0,002 vs PSACCs IN |
|  | PSACCs IN | **957,33 ± 184,98**<br>*p*<<0.001 vs IPSACs IN | **425,84 ± 60,86**<br>*p*<<0.001 vs IPSACs IN | **706 ± 93,28**<br>*p*<<0.001 vs IPSACs IN | **405,59 ± 53,81**<br>*p*<<0.001 vs IPSACs IN | **-14,15 ± 3,46**<br>*p*<<0.001 vs IPSACs IN | **869,11 ± 106,27**<br>*p*<<0.001 vs IPSACs IN |
|  | IPSACs PN | **7,51 ± 2,50**<br>*p*=0,021 vs IPSACs IN | 4,66 ± 1,21 | 12,56 ± 1,58 | 5,16 ± 0,78 | **-246,86 ± 68,18**<br>*p*=0,029 vs IPSACs IN | 15,38 ± 1,94 |
|  | IPSACs IN | 2,77 ± 1,00 | 5,74 ± 1,22 | 16,43 ± 3,09 | 5,94 ± 1,48 | -177,40 ± 84,93 | 20,62 ± 4,82 |

Cl^-^ conductances in neurons can be generated by activation of either GABAergic or glycinergic receptors. During glutamatergic blockade with 3 mM KYN, slice perfusion with 15 µM of the GABA_A_ receptor antagonist, gabazine, markedly reduced 4AP-induced PSACCs in 9 PNs (from 1023,76 ± 299,36 to 51,60 ± 21,71 pA) and in 8 INs (from 478,10 ± 112,08 to 43,04 ± 12,62 pA; Fig. 3a,b). The blocker of the GABA_A_-activated Cl^-^ ionophore, picrotoxin^21^ 100 µM (n=12), also reduced PSACCs amplitude (from 2808,55 ± 764,62 to 211,54 ± 95,50 pA in 7 PN and from 1517,81 ± 493,17 to 189,84 ± 68,05 pA in 5 IN; Fig. 3c,d). As illustrated in Figures 3a and 3c, both drugs also blocked the fast chloride currents (inhibitory postsynaptic current – IPSC), with a slower effect of PTX on PSACCs than on IPSCs. A similar effect was also observed when another GABA_A_ receptor antagonist, bicuculline (50 µM), was applied to slices (n=10; data not shown). Finally, the glycine receptor antagonist, strychnine (STR; 500 nM), did not affect amplitude of both currents (Fig. 3e,f); the rate of occurrence of PSACCs was reduced by STR (right panel in Fig. 3g, from 5 PNs and 4 INs). These experiments suggest that both the PSACCs and the fast IPSCs were mediated by the activation of the Cl^-^ ionophore associated with the GABA_A_ receptors.

**Figure 3.**
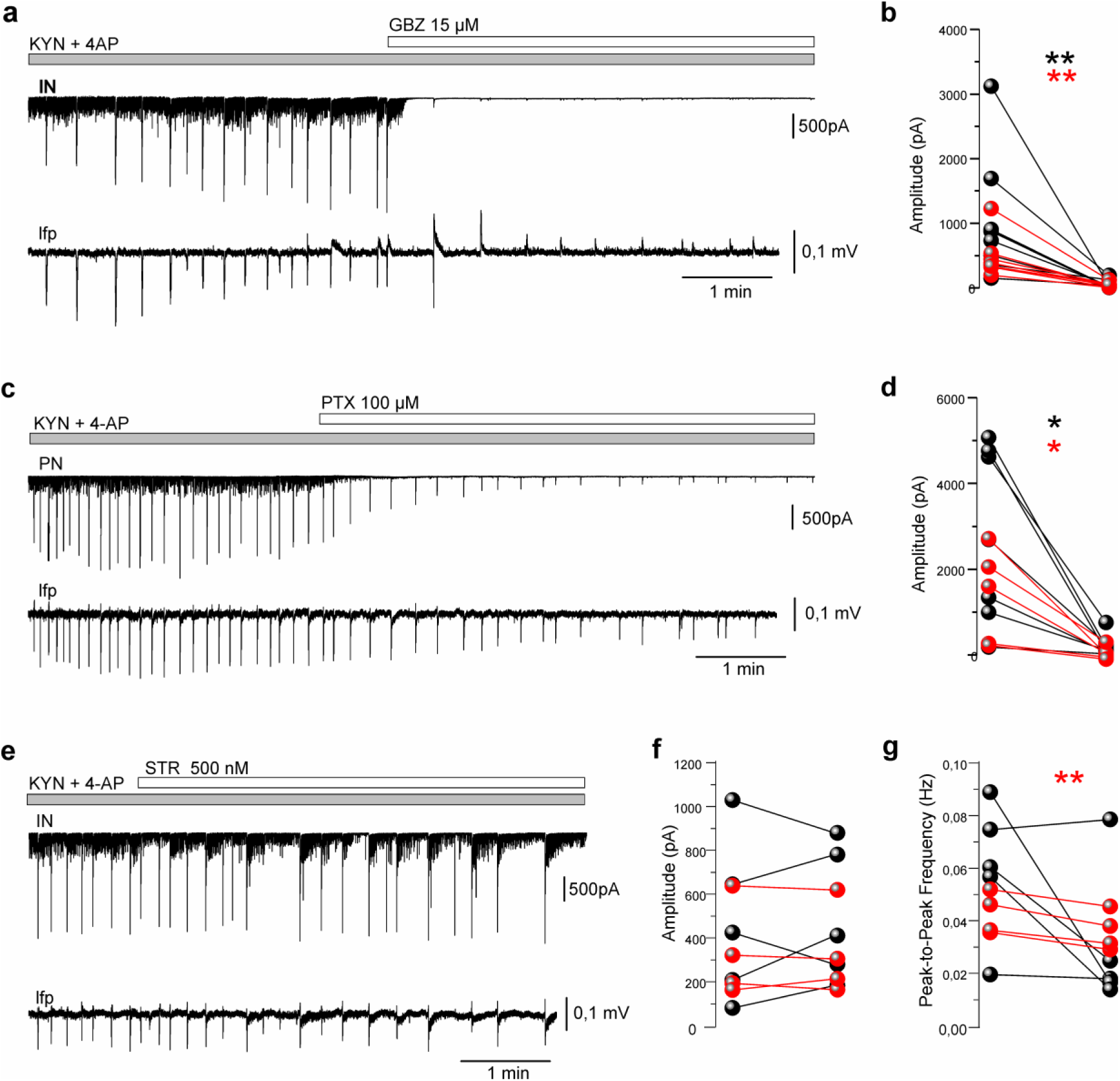
Effect of GABA_A_ and glycine receptor antagonists on PSACCs and IPSCs, during EC slice co-perfusion of 3 mM KYN and 100 µM 4AP (grey bars). **a.** Representative experiment performed with voltage-clamp intracellular recording (upper traces) and extracellular lfp (lower traces) before and after 15 µM gabazine (GBZ) application (white bar on top) is illustrated for an IN. **b.** PSACCs amplitude changes in 9 PNs (black symbols) and 8 INs (red symbols) before (left column of symbols) and after GBZ perfusion (right column). Values measured in the same neuron are connected by a line. **c.** Effect of 100 µM picrotoxin (PTX) perfusion recorded from a PN. **d.** PSACCs amplitude changes in 7 PNs (black) and 5 INs (red) before and after PTX perfusion. **e.** Effect of 500 nM strychnine (STR) perfusion in an IN. **f.** PSACCs amplitude changes in 5 PNs (black) and 4 INs (red) before and after STR perfusion. **g.** PSACCs frequency changes in PNs (black) and INs (red) before and after STR perfusion. Paired T-test and Wilcoxon Mann Whitney test were used. Statistical significance: * p<0.05, **p<0.01, *** p<0.005.

Next, we analysed the effect of synaptic transmission modulation on both IPSCs and PSACCs. Perfusion of the voltage gated sodium channel blocker, tetrodotoxin (1 µM; n=12), and the calcium channel blocker, cadmium chloride (CdCl_2_ 100 µM; n=11) in presence of 4AP (100 µM) and KYN (3 mM) abolished both slow and fast Cl^-^ currents in all cell types (from 768,18 ± 175,02 to 101,90 ± 44,51 pA in 6 PN and from 1042,45 ± 393,19 to 82,95 ± 39,93 pA in 6 IN; from 1396,76 ± 493,33 to 146,14 ± 66,44 pA in 6 PN and from 882,15 ± 206,17 to 131,07 ± 58,29 pA in 5 IN for TTX and CdCl_2_ respectively, as illustrated in Fig. 4a-d). Unexpectedly, the divalent cation cobalt chloride (CoCl_2_ 1 mM; n=20), did not significantly modify PSACCs amplitude (from 711,32 ± 193 to 586,48 ± 110,45 pA in 10 PN and from 786,60 ± 237,18 to 561,19 ± 148,58 pA in 10 IN; Fig. 4e,f), but reduced their recurrence rate in both INs and PNs (Fig. 4e,g). In 5 cells, an increase in PSACC amplitude was observed (see Fig. 4f and Discussion). Like CdCl_2_, CoCl_2_ consistently abolished IPSCs amplitude and frequency (Fig. 4e and triangles in Fig. 4f,g; n=10). Overall, this set of experiments suggests that GABA-mediated PSACCs generated by 4AP in the presence of glutamate receptors antagonist depends on the presence of synaptic neurotransmitters release.

**Figure 4.**
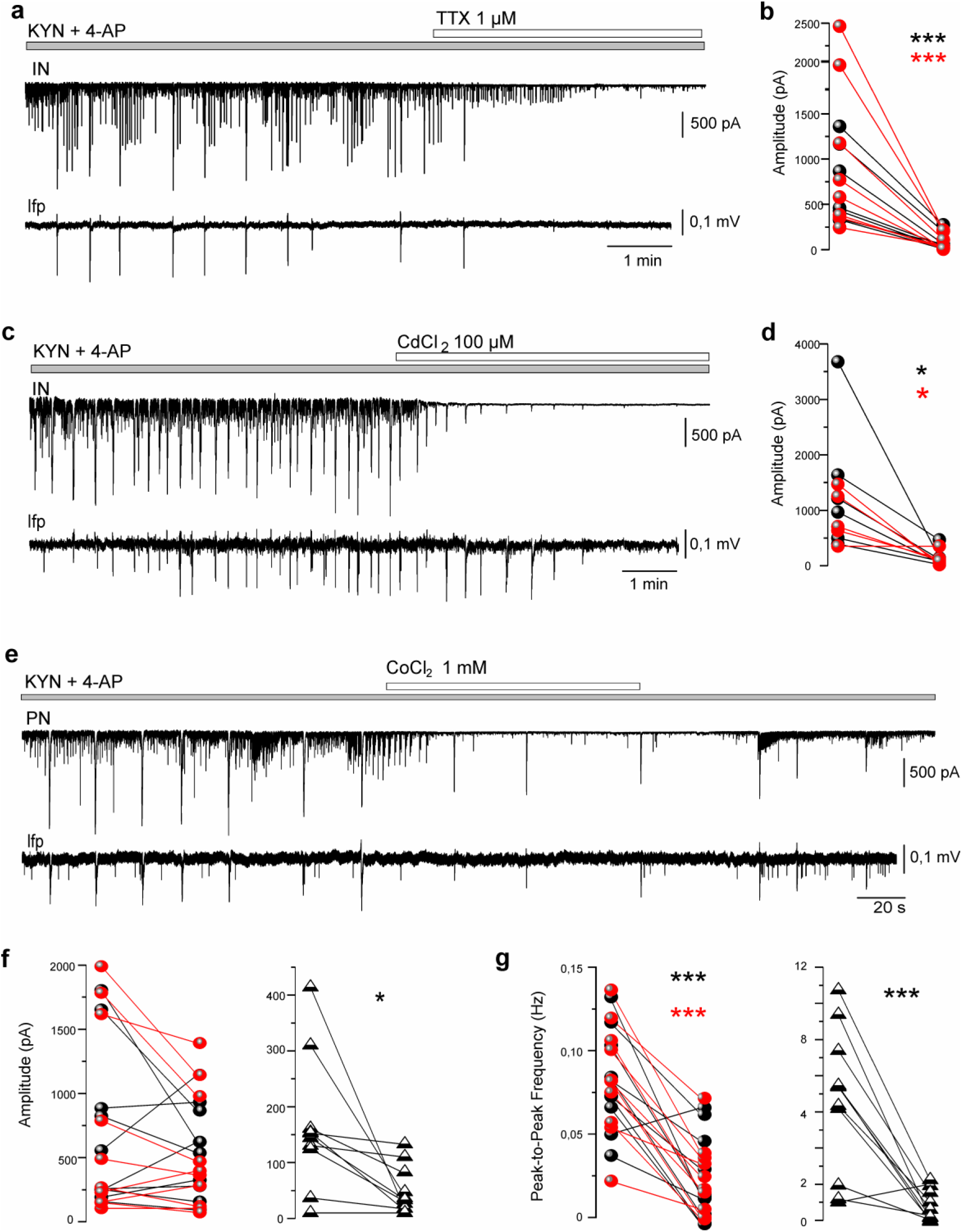
Effect of sodium and calcium channel blockers on PSACCs and IPSCs during EC slice co-perfusion with 3 mM KYN and 100 µM 4AP (grey bars). On the left, representative traces obtained with voltage-clamp intracellular recording (upper traces) and extracellular lfp (lower traces). On the right, pairs of current amplitude measurements performed before and after drugs application are illustrated for PNs (black spheres) and INs (red spheres). Values measured in the same neuron are connected by a line. **a.** Effect of 1µM tetrodotoxin (TTX; horizontal white bar on top) perfusion in an IN. **b.** PSACCs amplitude changes in PNs (n=6) and INs (n=6) before and during TTX perfusion. **c.** Effect of 100 µM cadmium chloride (CdCl_2_) recorded from an IN. **d.** PSACCs amplitude changes in PNs (n=6) and INs (n=5) before and after CdCl_2_ application. **e.** Effect of 1 mM cobalt chloride (CoCl_2_) on a PN, followed by CoCl_2_ washout. **f.** PSACCs (spheres) and IPSCs (triangles) amplitude changes in PNs (n=10) and INs (n=10) before and after CoCl_2_ perfusion. **g.** PSACCs and IPSCs frequency changes in PNs (n=10) and INs (n=10) before and after CoCl_2_ application. Paired T-test and Wilcoxon Mann Whitney test were used. Statistical significance: * p<0.05, **p<0.01, *** p<0.005.

### PSACCs and neurotransmitter transporters

Next, we analysed the effects of neurotransmitter transporters on PSACCs. EC slices perfusion with the excitatory amino acid transporter (EAAT) antagonist DL-threo-beta-benzyloxyaspartate (TBOA, 100 µM; n=12) reduced PSACCs amplitude in both PNs and INs (from 422,12 ± 82,89 to 213,90 ± 45,60 pA in 7 PNs and from 868,45 ± 321,04 to 607,14 ± 293,52 pA in 5 INs) and the associated population spikes; this effect was more consistent in PNs (black asterisks in Fig. 5b). TBOA also reduced recurrence rate from 0,09 ± 0,01 to 0,03 ± 0,009 Hz in 7 PNs and from 0,11 ± 0,02 to 0,02 ± 0,006 Hz in 5 INs (Fig. 5a-c). When the GABA transporter 1 antagonist, (NO-711; 50 µM; n=13) was added to the slices solution, the PSACCs recurrence rate was significantly reduced (from 0,05 ± 0,003 to 0,017 ± 0,001 Hz in 7 PNs and from 0,07 ± 0,01 to 0,02 ± 0,01 Hz in 6 INs; Fig. 5d,f) and PSACCs duration was markedly increased (from 433,74 ± 72,77 to 1429,39 ± 208,23 ms in 7 PNs and from 811,51 ± 183,70 to 2977,27 ± 790,78 ms in 5 INs; Fig. 5d,g). Interestingly, the occurrence of IPSCs increased after the occurrence of the enlarged PSACCs, as shown in the representative trace illustrated in Figure 5d (see discussion). Moreover, the amplitude of the associated population spikes increased, and their morphology changed in parallel with the enlargement of the PSACCs, suggesting a modification in the population spike reversal potential and confirming a close relationship between PSACCs and population spikes.

**Figure 5.**
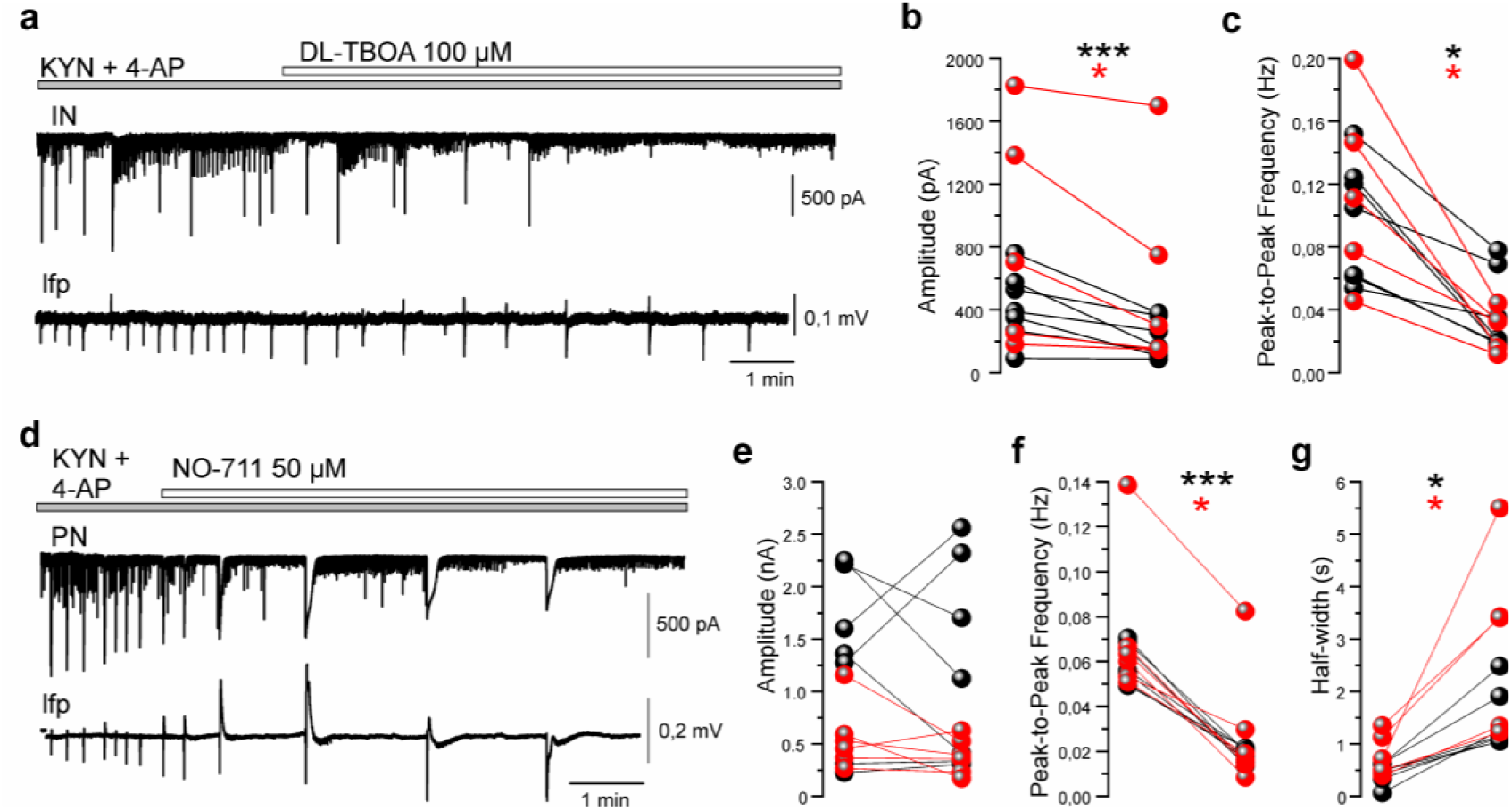
Effect of transporter antagonists on PSACCs and IPSCs during EC slice co-perfusion with 3 mM KYN and 100 µM 4AP (grey bar). On the left, representative traces of voltage-clamp intracellular recording (upper traces) and extracellular lfp (lower traces). **a.** Effect of 100 µM TBOA (white bar) in an IN and the relative lfp. **b.** Pairs of PSAACs amplitude measurements performed in PNs (n=7, black symbols) and INs (n=5, red symbols) before and after TBOA application. Values measured in the same neuron are connected by a line. **c.** PSACCs frequency measurements before and after TBOA application; PNs (n=7, black symbols) and INs (n=5, red symbols). **d.** Effect of GAT1 blocker, NO-711 (50 µM; white bar) in a PN. **e.** PSACCs amplitude values on PNs (n=7) and INs (n=6) before and after NO-711. **f.** NO-711 effect on PSACCs frequency recurrence; PNs (n=7) and INs (n=6). **g.** PSAACs duration, measured as changes in PSACCs half width, before and after NO-711; PNs (n=7) and INs (n=6). Paired T-test and Wilcoxon Mann Whitney test were used. Statistical significance: * p<0.05, **p<0.01, *** p<0.005.

Finally, PSACCs were dramatically reduced in amplitude in all recorded cells (from 2512,63 ± 414,06 to 626,23 ± 157,28 pA in 9 PNs and from 2022,02 ± 387,74 to 421,23± 168,77 pA in 5 INs) by 5-nitro-2-3-phenylpropylamino-benzoic acid (NPPB 100 µM; white bars in Fig. 6 a,c), a drug that antagonizes the Ca^2+^- activated bestrophin-1 (BEST-1) channel located mainly in astrocytes^22^ (as shown in Fig. 7), responsible for GABA release independent on synaptic vesicles. Population spikes were gradually abolished in parallel with PSACCs blockade (Fig. 6a,c). In 4 experiments, the perfusion of NPPB was preceded by the application of 1 mM CoCl_2_ (dark grey bar in Fig. 6c) to isolate PSACCs from synaptically released GABA; further addition of NPPB to the slice solution abolishes PSACCs (Fig. 6c). The experiments described above suggest that PSACCs i) are modulated by glutamate uptake via neurotransmitter transporters, ii) are enhanced by GABA transporter blockade and iii) are suppressed by blocking GABA release mediated by BEST-1 channels.

**Figure 6.**
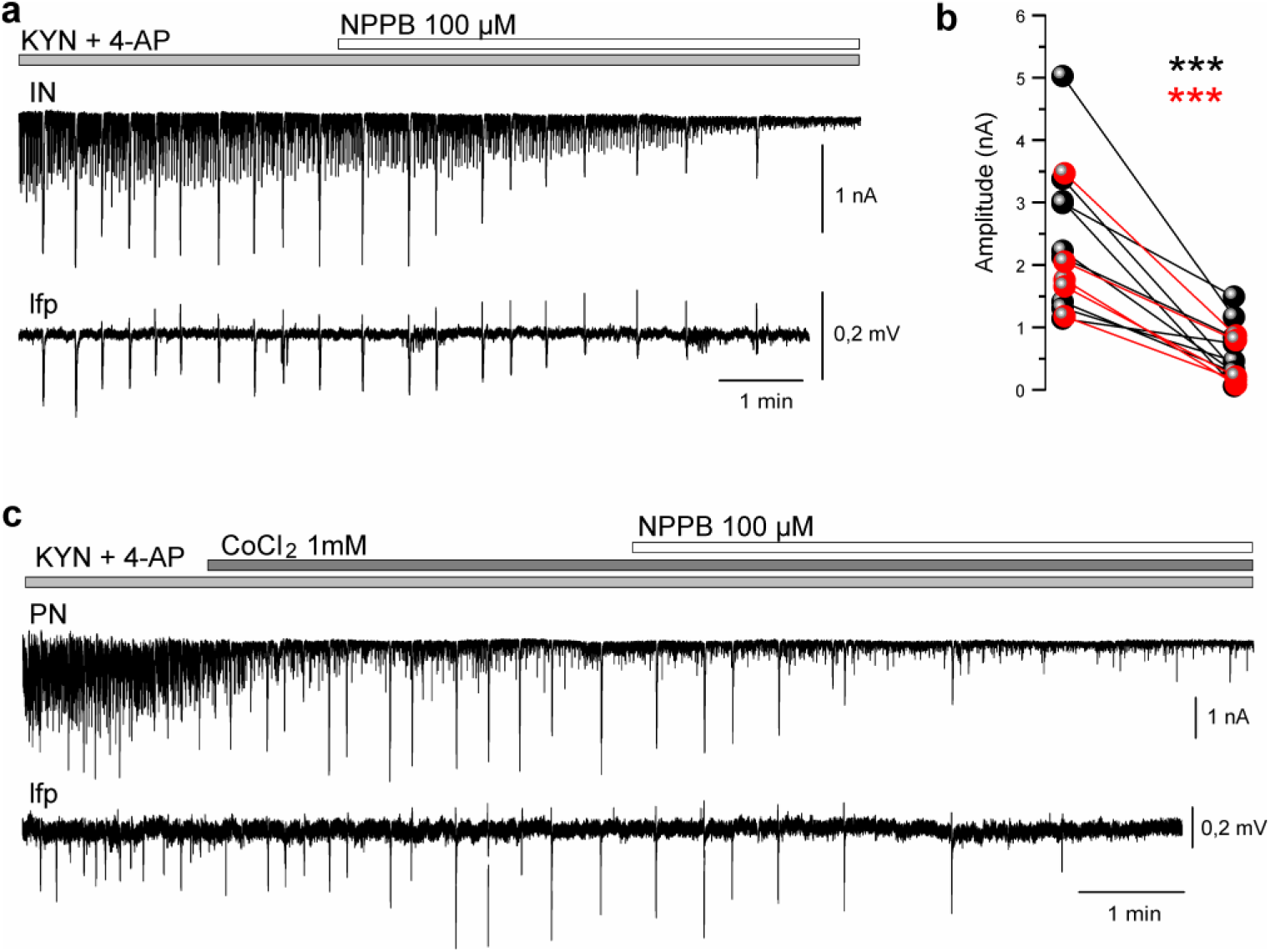
Effect of bestrophin 1 blocker on PSACCs and IPSCs under EC slices co-perfusion with 3 mM KYN and 100 µM 4AP (light grey bars). **a.** Representative voltage-clamp intracellular recording (upper trace) and extracellular lfp (lower trace) during application of 100 µM 5-nitro-2-3-phenylpropylamino-benzoic acid (NPPB; white horizontal bar). **b.** Pairs of amplitude measurements performed before and after NPPB are illustrated for 9 PNs and 5 INs.**c.** Similar experiment as in **a**, but in presence of CoCl_2_1 mM (dark grey bar on top). Remarkably, CoCl_2_ had no effect on PSACC, while it completely abolished spontaneous IPSC activity. Addition of NPPB abolished PSACCs. Paired T-test and Wilcoxon Mann Whitney test were used. Statistical significance: * p<0.05, **p<0.01, *** p<0.005.

**Figure 7.**
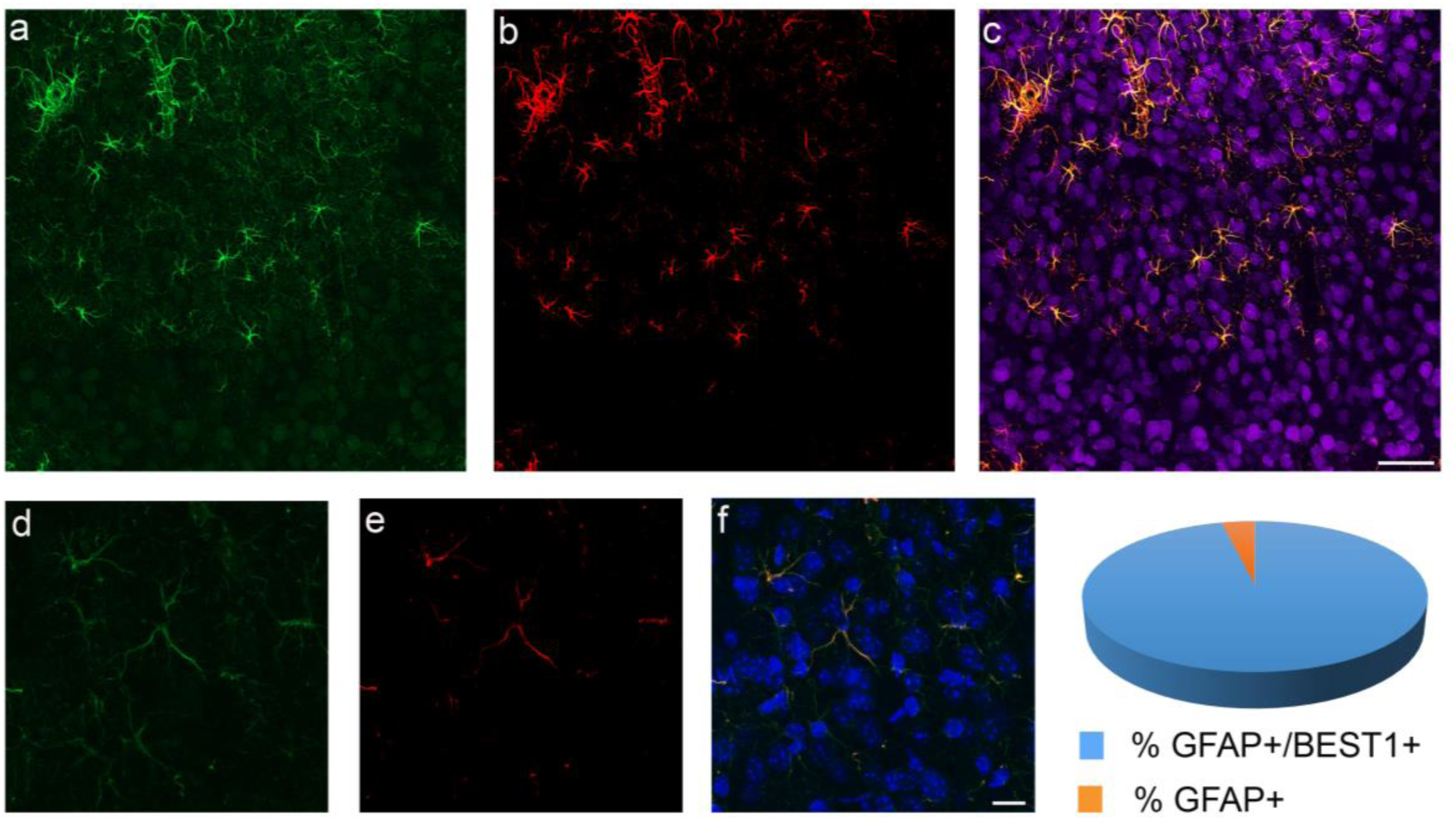
Bestrophin 1 co-localization with GFAP in mouse EC. Representative 20x confocal *z*-stack image showing triple immunofluorescence for (**a**) GFAP (in green), (**b**) BEST-1 (in red), and (**c**) merged BEST1/GFAP/NeuN (in pink). BEST-1 co-localise with GFAP positive cells and is not detectable in neurons. **d-e-f.** High magnification of a BEST1/GFAP; nuclei are counterstained with DAPI. Scale bars equal 50 μm in a-c; 10 μm in d-f. **g**. Pie chart showing the percentage of astrocytes (GFAP^+^ cells) expressing BEST1. Data were obtained from 112 GFAP positive cells counted in 8 EC slices (70 µm thick) cut from 2 mouse brains.

### Effect of BEST-1 blocker on SLEs

After dissecting the effects of different agents on 4AP-induced PSACCs, we verified the competence of NPPB to interfere with SLEs generated in EC slices in the presence of glutamate transmission^18^. Typical SLEs induced by 100 µM 4AP (top grey bar) are illustrated in Figure 8a (arrows mark SLE onset). When 4AP was co-perfused with 100 µM NPPB, SLEs were abolished (n=9; Fig. 8b) and were substituted by enhanced synaptic activity and population spikes. Unlike PSACC-mediated spikes observed with glutamate receptor blockade (Fig. 2c), the population spikes observed during NPPB were faster and were not consistently correlated with neuronal activity (n=7; see insert in Fig. 8b).

**Figure 8.**
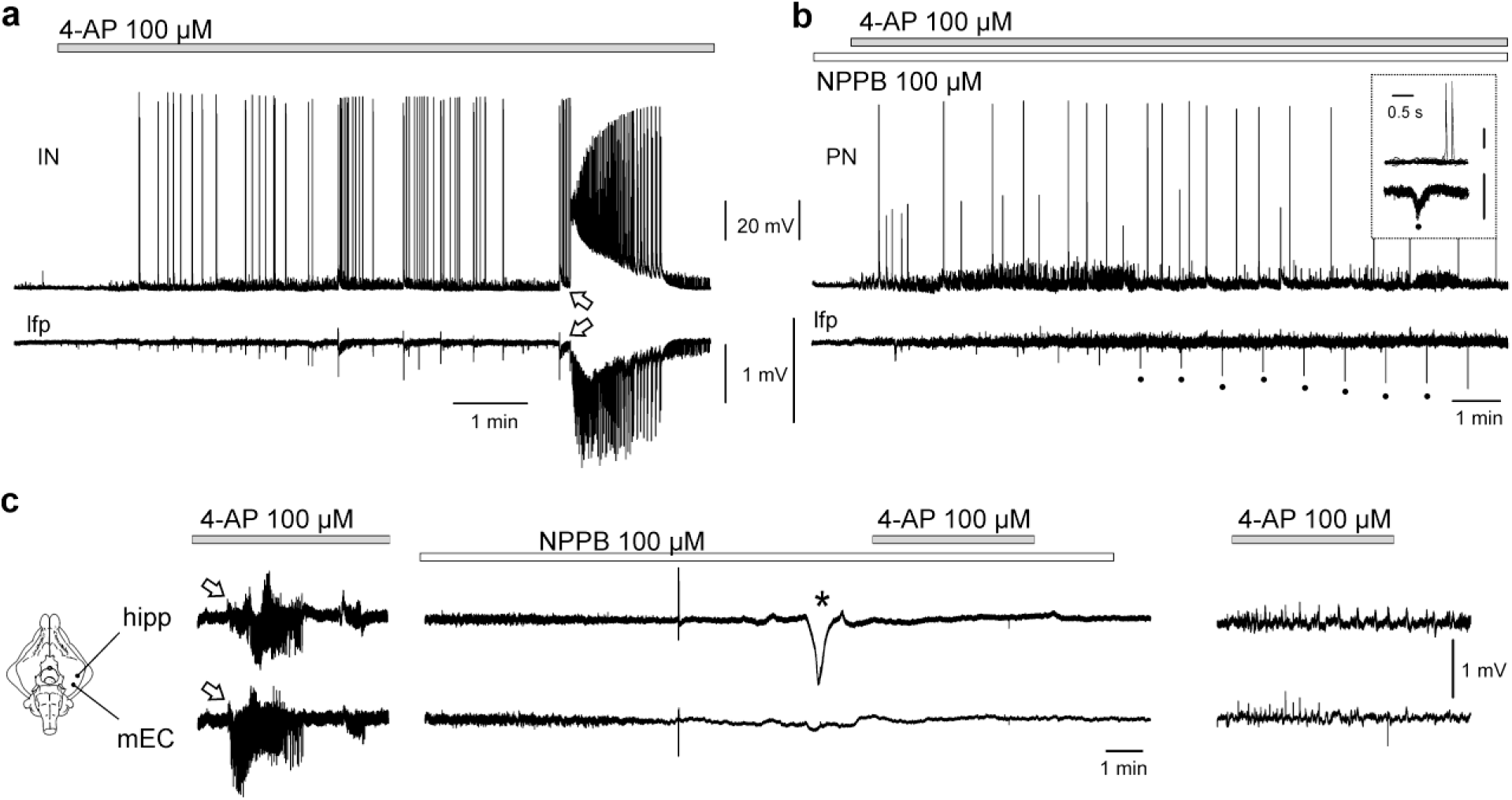
**Effects of PSACCs–active drugs on the generation of seizure-like events (SLEs) induced by 4AP in EC slices (a-d) and in the *in vitro* isolated guinea pig brain (e-f).** a. Current clamp recording of an IN (upper trace) and the simultaneous lfp (lower trace) during 100 µM 4AP perfusion, show the generation of a seizure-like event (SLE; arrow). **b.** Perfusion of the BEST-1 antagonist NPPB (white line) during 4AP (grey line) prevents the generation of SLE in a PN. In the insert superimposed population spikes (lower trace) and the associated neuronal activity (upper trace). **c.** Simultaneous extracellular recordings from the CA1 region of the hippocampus (hipp) and the mesial entorhinal cortex (mEC) of the *in vitro* isolated guinea pig brain (scheme on the left) during intra-arterial perfusion of 100 µM 4AP (grey bar) before, during and after intra-arterial perfusion of BEST-1 blocker NPPB (100 µM; white horizontal bar). A SLE is recorded before NPPB perfusion (open arrows) and epileptiform activity was observed after 1.5-2 hours NPPB washout (right part of the panel); during the perfusion with the BEST-1 antagonist no seizures were observed. A brief spreading depolarization during NPPB is marked by the asterisk.

Finally, to extend the observation obtained in mouse EC slices to a close-to-*in vivo* condition, we evaluated the effect of NPPB on SLEs induced by 5-8 minute arterial perfusion of 100 µM 4AP in the limbic cortices of the *in vitro* isolated guinea pig brain preparation^23,24^. When 100 µM NPPB was perfused for 15 minutes and then co-perfused with 4AP (5-8 min), SLEs generation was prevented in both the mesial EC (mEC) and the CA1 region of the hippocampus (hipp; middle panel in Fig. 8c; n = 5); in 3 experiments, NPPB alone (before 4AP co-perfusion) determined brief spreading depolarization-like events (asterisk in Fig. 8c) that originated in the hippocampal region and could propagate to the mEC. In 3 out of 3 NPPB experiments, after 1.5-2h of washout, re-application of 4AP for 5-8 minutes activated epileptiform interictal discharges but did not induce SLEs (as in the representative experiment illustrated in the right panel of Fig 8c), suggesting a long lasting effect of the BEST-1 blocker.

## Discussion

We demonstrate for the first time that the interictal population spikes induced by 4AP in mouse EC slices are closely coupled with a large and synchronous extrasynaptic GABA-mediated chloride conductance in all recorded neurons specifically identified as PNs or INs by soma features, firing properties and GAD65/67 fluorescent reporter^19^. The PSACCs has unambiguously different kinetic properties compared to the fast IPSCs mediated by synaptic GABA_A_ receptors: PSACCs last > 2 sec, peak at 300-1000 ms (< 50 ms for IPSCs), decay in 400-1200 ms (*<*100 ms for IPSCs) and had large charge area (300-2500 nAmp/ms *vs* <100 nAmp/ms in IPSCs). Like IPSCs, PSACCs were sensitive to GABA_A_ receptor antagonists and were abolished by neurotransmitter release blockers. Unlike IPSCs, PSACCs i) were not blocked by CoCl_2_, ii) were reduced by EAAT antagonist, iii) were augmented by GAT1 antagonist, iv) were suppressed by blocking BEST-1 channels.

PSACCs are not generated by glutamatergic excitatory transmission, since they persisted in the presence of glutamate receptor antagonists. The effect of gabazine, bicuculline and GABA_A_ receptor associated chloride channel blocker, picrotoxin^21^, confirmed that PSACCs are mediated by GABA, acting on GABA_A_ receptors. As described for the GABA-mediated Cl^-^ conductance recorded in slices with 19 mM chloride intra-pipette solution^25^, PSACCs reversal potential was -50 mV and shifted toward more depolarized values when the intracellular Cl^-^ was increased by raising its concentration in the patch pipette. Since PSACC was specifically abolished by BEST-1 channel blocker, we propose that it is mediated by GABA released from BEST-1 channels through a non-synaptic mechanism via extrasynaptic GABA_A_ receptors (see below).

PSACCs blockade by CdCl_2_, could be explained by several possible action: Cd^2+^ influences GABA_A_ receptor activation^26,27^ and it is responsible for the inhibition of GABA transporters^28^. CdCl_2_ could be responsible for BEST-1 channels blockade due to its competition with intracellular Ca^2+^ binding sites that impair channel opening. Interestingly, the other divalent cation (CoCl_2_ 1 mM) utilized in our experiments abolishes synaptic GABAergic transmission (IPSCs) without affecting PSACCs amplitude, which increases in some experiment. This phenomenon could be due to a greater availability of free post synaptic GABAergic receptors due to the blockade of synaptic activity by cobalt, that over time reduces PSACC frequency, as shown in Figure 4g. There are no data in literature on the effect of CoCl_2_ on GABA transporters in mammalians brain. These findings demonstrated that PSACCs were GABA_A_ receptor-mediated Cl^-^ currents due to a non-vesicular mechanism that does not depend on synaptic GABA release.

The experiments with blockers of neurotransmitter transporters demonstrated that PSACCs depend on the enhanced synaptic release of GABA and/or glutamate promoted by 4AP, a potassium channel blocker that prolongs the duration and frequency of the presynaptic action potentials^29,30^. The released glutamate is removed from the synaptic cleft through a transmembrane exchange with Na^+^ mediated by EAATs, in particular by EAAT1 expressed on astrocytic processes^31,32^ in close vicinity to the glutamatergic axon terminals^33–35^. Glutamate uptake by astrocytes via EAAT is, indeed, necessary for the induction of activity-dependent astrocytic responses^36^. In our experiments, the EAAT antagonist, TBOA, reduced both PSACCs amplitude and frequency, suggesting that the reuptake of glutamate is required to generate PSACCs. Interestingly, PSACCs were also sensitive to TTX, possibly because of its action on presynaptic neuronal firing responsible for the accumulation of glutamate (and also GABA) that is transported into cells by EAATs, since postsynaptic glutamate receptors were blocked by the action of KYN acid. The TTX blocking effect confirms that neuronal firing is essential for neuron-astrocyte communication^36^ and that TTX impairs gap-junctional communication between neurons and astrocytes^37^. Moreover, the blocker of GAT1 (NO-711) increased PSACCs duration and the overall ion charge carried by the current. Interestingly, this effect was coupled with an increased IPSCs frequency just after each PSACC (Fig. 5d), suggesting a spread of GABA released during PSACCs from the peri-synaptic region into the synaptic cleft. The failure of GABA reuptake due to the blockade of GAT1 caused an increase of extracellular GABA concentration, which likely reflects the changes in amplitude and inversion of the dipole responsible for the lfp spike. A similar effect possibly due to enhanced GABA availability could mediate the changes in extracellular spikes transiently observed during GBZ perfusion.

Recent evidence in animal models of epilepsy suggests that the tonic extra-synaptic GABAergic inhibition could be due to GABA synthesis and release by reactive astrocytes^38–40^. Cultured astrocytes release GABA when activated by kainic acid stimulation and this process is blocked when Na^+^ is removed from the extracellular solution^41^. Interestingly, glutamate application in rodent cortical slices induced radiolabeled GABA release that was independent on both glutamate receptor antagonist and Ca^2+^-mediated vesicular release but relied on glial neurotransmitter transporters GAT and EAAT^34^. Extra-synaptic GABA could be released from astrocytes either via GABA transporters operating in the reverse mode (as demonstrated in Bergman glia of cerebellum^42,43^) or by permeation through the BEST-1 anion channel specifically expressed in glia cells^38,44^. The most interesting and original finding of our study is the demonstration that BEST-1 antagonist NPPB, consistently caused PSACCs amplitude reduction. BEST-1 channels are present in the brain, mainly on astrocyte membranes^40,45^ as confirmed by our data, and mediate non synaptic release of glutamate and GABA^22,38,40^, either by a Ca^2+^ dependent mechanism^22,46–50^ and by cellular swelling^47^. Interestingly, seizure activity coupled with a reduction in hippocampal pyramidal cells tonic GABA inhibition was observed in *Best1* gene knock out mice^51^. In this model, the overexpression of *Best1* specifically targeted to hippocampal astrocytes restored tonic GABA currents and decreased seizure recurrence^51^. Finally, BEST-1 channels were directly activated not only by intracellular Ca^2+^, but also by GABA through a direct GABA binding on the channel extracellular sites, suggesting that BEST-1 is controlled by ligands from both sides of the membrane^52^. This study also demonstrated that BEST-1 GABA-activated Cl^-^ channel shares no structural homology with GABA_A_ receptors. Based on these premises, we hypothesize that PSACCs are generated in both glutamatergic (PNs) and GABAergic neurons (INs) by GABA released via astrocytes BEST-1 channels in proximity to synapses.

The PSACC is similar to previously observed slow events attributed to glia release of neurotransmitters^53–55^. Astrocytic release of glutamate during epileptiform discharge was demonstrated in hippocampal slices and *in vivo* during application of 4AP and bicuculline^56^. Two-photon Ca^2+^ imaging combined with field recordings showed that Ca^2+^ waves in astrocytes induced either by convulsants and by caged Ca^2+^ photolysis preceded the onset of population epileptiform spikes^56^. Similar effects were observed with electrical stimulation in hippocampal mouse slices in the presence of 4AP^36^. Unlike PSACCs, the slow inward currents due to glutamate astrocyte release in the presence of glutamatergic transmission reversed at about 0 mV and were blocked by glutamatergic NMDA receptor antagonists in hippocampal slices^56–58^; these glutamate-driven events were not abolished by TTX and were enhanced by glutamate transporters inhibitor, TBOA^57^. Moreover, slow outward currents generated by GABA released from astrocytes were demonstrated in olfactory bulb slices^59^, in hippocampal slices in response to mechanical astrocytes stimulation and in hypo-osmotic solutions^60^ and in layers 2/3 neurogliaform cells of the rat somatosensory cortex^61^. GABA gliotransmission in the mammalian brain was extensively reviewed^62^, but its role in the framework of epileptic ictogenesis is not defined yet. Interestingly, a recent report demonstrated rhythmic release of GABA during interictal activity in mouse hippocampal slices perfused with a low Mg^2+^/high K^+^ solution^63^. As in our model, the extracellular GABA increase indirectly measured by recording Cl^-^ currents with single channel patch pipette (GABA sniffer) correlated to the generation of interictal population spikes in the field recording.

Based on the above summarized data, we developed a model of the sequence of events responsible for PSACCs generation in the mouse EC bathed in 4AP; the model includes an excitatory synapse (on the left in Fig. 9), a GABAergic synapse (on the right) and an astrocyte process (in the middle). During physiological neuronal activity, astrocytes are responsible for clearing the extracellular space from the excess of neurotransmitters^64^ (*phase 1* in Fig. 9). During 4AP application, presynaptic action potentials are prolonged, and more neurotransmitter is released at both glutamatergic and GABAergic synapses (*phase 2* in Fig. 9). The glutamate is taken up by astrocytes and is transformed in GABA by glutamic acid decarboxylase^62,65^ and accumulates in the astrocytic processes adjacent to synapses. GABA accumulation in distal astrocytic processes may generate cellular swelling that promotes periodic and synchronous Ca^2+^-mediated GABA release by astrocyte BEST-1 channels^50,60^ (*phase 3* in Fig. 9). This massive release of GABA induces the observed large PSACCs in postsynaptic GABAergic synapses, possibly by acting via extrasynaptic GABA_A_ receptors (*phase 3* in Fig. 9).

**Figure 9:**
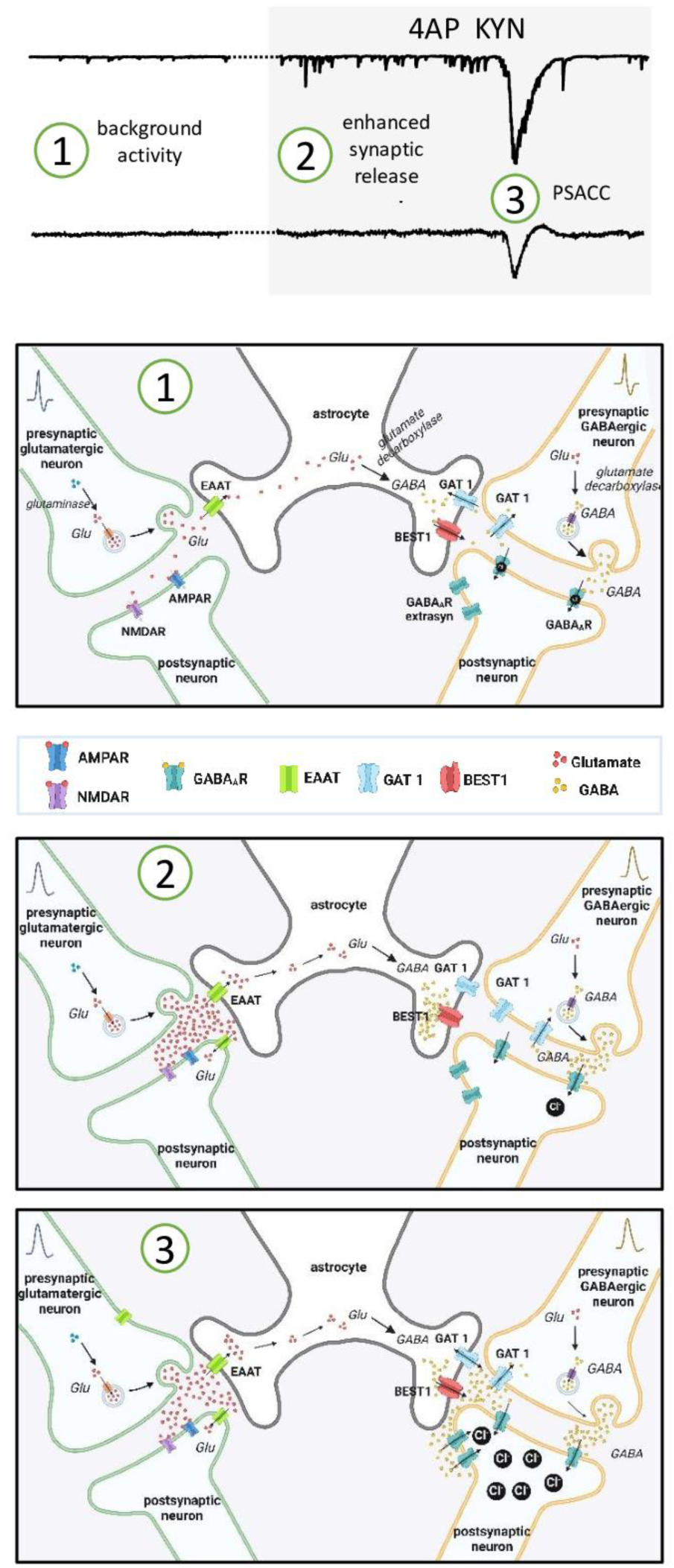
**Scheme of EC focal ictogenesis in the 4AP seizure model**. The trace in the upper panel shows a typical PSACC (3) recorded in EC mouse slices, preceded by enhanced synaptic activities (2) observed during coperfusion of 4AP and kynurenic acid (KYN); the background activity recorded with control solutions (1). The schemes (created with **BioRender.com**) illustrate the proposed mechanisms associated with the 3 phases highlighted in the traces. The schematic drawing includes an excitatory synaptic terminal (left) a GABAergic terminal (right) and an astrocyte termination on both synapses (center). The legend of the different symbols is reported at the bottom of the figure.

The large Cl^-^ influx associated with the PSACCs is likely re-balanced by the immediate activation of K^+^/Cl^-^ co-transporter KCC2 that leads to the extracellular K^+^ accumulation^66,67^ observed during population spikes just ahead of a seizure event in the 4AP model^14,18^. The massive release of GABA during PSACCs could also activate GABA_B_ metabotropic receptors, causing GIRK activation and the related outflow of K^+^ ^68^. As mentioned above, the increase in extracellular K^+^ concentration could cause the initiation of the epileptic seizure. In addition, the large and fast intracellular Cl^-^ increase during PSACCs could *per se* promote seizures^69,70^; high intracellular Cl^-^ concentration is known to facilitate neuronal firing and to promote seizures by decreasing the threshold for action potential generation^71^. Finally, Cl^-^ accumulation in neurons causes a positive shift in the polarity of the electrochemical Cl^−^ gradient that results in E_GABA_ depolarizing shift above the action potential threshold; in these conditions, ion channel opening will lead to an outward flow of Cl^−^ that will sustain membrane depolarization, and possibly seizures.

The close correlation between the population spikes and the PSACCs during pharmacological manipulations has demonstrated that these two events represent the same phenomenon observed from different recording viewpoints: intracellular *vs* lfp. Parallel variations between PSACCs and spikes during the pharmacological tests performed in our study, strongly support the idea that 4AP-induced spikes are generated by the mechanisms supported by PSACCs in the absence of glutamatergic transmission due to KYN. When excitatory synapses are not blocked, the synchronous presence of spikes and PSACCs initiate SLEs. In line with this, we demonstrated that BEST-1 channel blocker prevents the occurrence of SLEs in EC slices perfused with 4AP in full preservation of both excitatory and inhibitory synaptic transmission. NPPB was also interfering with 4AP-induced SLEs in a close-to-*in vivo* isolated guinea pig brain. These findings confirmed that the interference with mechanisms underlying PSACCs can control ictogenesis in the 4AP focal seizure model.

### Limitations of the study

*In vitro* reproduction of epileptiform discharges is a recognized and accepted surrogate of focal ictogenesis. 4AP application represents the most utilized *in vitro* model to reproduce interictal activity and SLEs that are similar to the patterns observed in human temporal lobe seizures^14,15^. Recordings of extracellular units can be performed *in vivo* to analyse the contribute of neurons during seizures. Still, the pharmacological dissection of the possible mechanisms that contribute to the generation of epileptiform discharges is limited in the *in vivo* condition by the side effects of the drugs utilized to block specific functions – such as BEST-1 antagonist. Moreover, even though the sequence of events here described are specific to the mechanism of action of 4AP, rhythmic GABA increases coupled with epileptiform spikes were also observed in the zero Mg^2+^/high K^+^ *in vitro* model^63^, suggesting that the mechanism here described could be suitable for different epileptogenic conditions.

## Conclusions

GABAergic spikes mediated by interneuronal activity have been described in the 90’s by Massimo Avoli group^11,12^ and were more recently confirmed in several *in vitro* temporal lobe seizure models^13,18^. Based on these early findings, the role of GABA and GABAergic networks in the generation of SLEs have been recognized and firmly demonstrated in different *in vitro* preparations, in animal models *in vivo* and in single unit recordings performed in patients with drug-resistant focal epilepsies during presurgical intracerebral monitoring^10,14,15^. However, the activation of excitatory neurons and glutamatergic drive in seizure generation is also well established in specific models of seizure and epilepsy^1,72^. Our findings demonstrate **a novel mechanism of ictogenesis** that depends on massive GABA release from astrocytes, which generates large Cl^-^ currents in all types of neurons, regardless of their competence to release glutamate or GABA as neurotransmitter. The demonstration of a GABA-driven Cl^-^ current in strict correlation with interictal and preictal spikes in the 4AP seizure model provides a new pathogenic function in seizure genesis for astrocyte-mediated GABA release; PSACC-associated postsynaptic Cl^-^ influx on a large number of neurons may lead to ion exchanges that promotes seizure initiation. The proposed PSACCs-mediated mechanism of ictogenesis obscures the specific role of excitatory and inhibitory synapses in seizure generation and supports the hypothesis that neurons are dragged into epileptiform discharges by hastily following the fast transmembrane ion changes imposed by non-vesicular GABA release sustained by astrocytes.

## Methods

Experiments were carried out on C57BL/6J mice (Charles River, Calco, Italy) according to the European directive 2010/63/UE, approved by Institutional Committee on Animal Care and Use and by National ethical committees (Protocol 546/2023-PR for brain slices and DO-01-2023 for the isolated guinea pig brain). ARRIVE guidelines and the Basel declaration were considered when planning the experiments. All efforts were made to minimize the number of animals and their suffering. Mice were group housed (5 per cage) on a 12 h light/dark cycle, with *ad libitum* access to water and food. Every effort was made to limit the number of animals used.

### Entorhinal cortex brain slices

Brain slices were prepared as previously described^73,74^ from 20 to 30 day-old C57BL/6J mice that specifically express the green fluorescent protein (GFP) in GABAergic interneurons (IN): GAD65-GFP transgenic mice^75^ and GAD67-GFPΔneo knock-in mice^76^. GAD67 knock-in mice were provided by Y. Yanagawa (Gunma University, Japan), GAD65 transgenic mice were provided by G. Szabo (Institute of Experimental Medicine, Budapest, Hungary). After mouse decapitation under isoflurane anesthesia (Isoflo, Zoetis Italia), the brain was quickly isolated and placed in ice-cold artificial solution containing (mM): 87 NaCl, 7 MgCl_2_, 2.5 KCl, 0.5 CaCl_2_, 21 NaHCO_3_, 1.25 NaH_2_PO_4_, 25 glucose and 75 sucrose, bubbled with 95% O_2_ 5% CO_2_. Horizontal slices (400 μm thick) that contained the entorhinal cortex (EC) were cut in the above ice-cold solution with a vibratome (Leica VT1200S; Wetzlar, Germany) and were placed in an incubation chamber bathed in a standard solution, which contained (mM): 129 NaCl, 1.8 MgSO_4_, 3 KCl, 1.6 CaCl_2_, 21 NaHCO_3_, 1.25 NaH_2_PO_4_ and 10 glucose (ACSF). Experiments were performed at room temperature.

### Isolated guinea pig brain in vitro

Young adult guinea pigs (150–250 g) were anaesthetized with with isoflurane (Isoflo, Zoetis Italia). During anaesthesia, intracardiac perfusion with cold (10°C) oxygenated saline solution (see below) was performed to reduce brain temperature. Brains carefully dissected out according to the standard technique^23,24^ were transferred to a recording chamber and were perfused via a peristaltic pump at 5.5 mL ⁄ min with a complex saline solution (composition in mM: 126 NaCl, 3 KCl, 1.2 KH_2_PO_4_, 1.3 MgSO_4_, 2.4 CaCl_2_, 26 NaHCO_3_, 15 glucose, 2.1 HEPES, and 3% Hydroxyethyl starch; Voluven M.W. 130 KDa; Fresenius Kabi, Isola della Scala, Italy) through a polyethylene cannula inserted into the basilar artery. The solution was saturated with 95% O_2_ ⁄ 5% CO_2_ (pH 7.3). During the surgical procedure the chamber temperature was slowly raised (0.2°C ⁄ min) to 32° C before starting the recordings. SLEs were induced by arterial perfusion of 100 µM 4AP for 5-8 minutes.

### Electrophysiological recordings

Electrophysiological recordings on *in vitro* slices started after an incubation/recovery period of at least one hour. Slices were transferred in the recordings chamber (Warner Instruments, Hamden, CT, US) and cells were visualized by infrared video microscopy with a Nikon Eclipse FN1 microscope (Nikon, Minato, Japan) equipped with DIC optics and a CCD camera (Hamamatsu, Japan). Simultaneous extracellular local field potentials (lfp) and whole-cell current-clamp patch-clamp recordings were performed at 25°C with Multiclamp 700B patch-clamp amplifier, Digidata 1440a digitizer and pClamp 10.2 software (Axon Instruments-Molecular Devices, San Jose, CA, US). Micropipettes for patch clamp and lfp recordings were pulled from borosilicate glass capillaries (Sutter Instruments, Novato, CA, US; input resistance 2.5-3.0 MΩ; access resistance of 4-7 MΩ). ACSF solution was used as lfp pipette internal solutions; internal solution for current-clamp patch-clamp recordings contained (in mM) 120 K-gluconate, 15 KCl, 2 MgCl_2_, 0.2 EGTA, 10 HEPES, 4 Na_2_ATP, 10 P-creatine, adjusted to pH 7.2 with KOH^19^. High Cl^-^ solution for voltage-clamp recordings contained (in mM) 135 CsCl, 2 MgCl_2_, 0.2 EGTA, 10 HEPES, 1,5 QX314 at pH 7.2 corrected with CsOH. Lfp recordings were performed with Multiclamp 700B amplifier in I=0 mode. Whole-cell patch-clamp recordings were performed from the cell soma; IN were identified by GFP fluorescence, PN were identified by their size and the pear-like soma and by the lack of GFP fluorescence^19^. Signals were filtered at 10 kHz and were sampled at 50 kHz. Signal analyses were performed using pClamp 10.5 and Origin 2021 (Origin Lab, Northampton, MA, US). To characterize neuron subtypes, input-output curves were obtained by injecting 2.5-sec depolarizing current pulses increasing by 10 mV steps. Action potential firing adaptation was analyzed during depolarizing current ramps ranging between 2 and 5 mV above firing threshold.

Lfp recordings in the CA1 area of the hippocampus and in the mesial EC (mEC) of the *in vitro* isolated guinea pig brains were performed with 0.9 M NaCl^-^-filled glass pipettes (5-10 MΩ resistance). Electrodes were inserted under stereoscopic microscope control and the position of the CA1 recording was achieved by analyzing the lfp evoked by electrical stimulation of the lateral olfactory tract^77^.

### Drugs

1,2,5,6-tetrahydro-1-2-diphenylmethyleneamino-oxyethyl-3-pyridinecarboxylic acid (NO-711); 4-aminopyridine (4AP); 5-nitro-2-3-phenylpropylaminobenzoic acid (NPPB); DL-threo-beta-benzyloxyaspartate (DL-TBOA); abazine (GBZ); kinurenic acid (KYN); picrotoxin (PTX); tetrodotoxin (TTX); strychnine (STR); cobalt chloride (CoCl_2_); cadmium chloride (CdCl_2_) and bicuculline were purchased at Sigma Aldrich and Tocris.

### Immunohistochemical procedures

Male C57BL6 wild-type mice were deeply anesthetized and transcardiacally perfused with 1% paraformaldehyde (PFA) in 0.1 M phosphate buffer (pH 7.2) followed by 4% PFA in the same buffer. Mouse brains were removed from the skull, immersed in 4% PFA overnight at 4°C and cutted into 50-μm thick serial horizontal sections by means of a Vibratome VT1000S (Leica, Heidelberg, Germany). Triple immunofluorescence was performed by incubating free-floating sections of mouse brains for 1 h in 10% normal goat serum in 0.01 M phosphate buffer saline (pH 7.4) and then in primary antibodies polyclonal rabbit anti-Best1 (1:400 Abcam, Cambridge, UK) in conjunction with a monoclonal mouse anti-GFAP (Millipore Merck, Darmstadt, Germany) and polyclonal guinea pig anti-NeuN (Millipore 1:3000 Millipore Merck, Darmstadt, Germany) overnight at 4°C. After a complete wash in phosphate buffer solution, sections were incubated in the secondary antibodies for 2 h: CY2–conjugated goat anti-mouse, CY3– conjugated goat anti-rabbit (1∶600; Jackson ImmunoResearch Laboratories, West Grove, PA, USA) and biotinylated goat anti-guinea pig (1:200 Vector Laboratories, Inc.Newark, CA, USA) followed by Alexa 647– conjugated streptavidin (1:600 Thermo Scientific, Rockford, IL, USA). Sections were counterstained with DAPI (1:1000 Thermo Scientific, Rockford,IL, USA) mounted with Fluorsave (Calbiochem, San Diego, CA, USA); EC was examined through a confocal microscope ZEISS LSM980 Confocal (Carl Zeiss AG, Germany) with a 20x or 63× objective. Automated sequential acquisition of multiple channels was used to obtain *z*-stack images.

### Statistics

Statistical analysis was performed using Prism 6.0 software (GraphPad, San Diego, CA, US) or Origin 2021 (OriginLab); T-test and Wilcoxon Mann Whitney test were used for single population data distributed either normally or not, respectively. For amplitude and frequency plots (Figs. 3-6), each value represents the average of 10 values before and 5 values after drug perfusion in each cell. Statistical significance: * p<0.05, **p<0.01, *** p<0.005.

## Acknowledgements

The study was supported by Italian Health Ministry PNRR-MAD-2022-12376068 grant and *Ricerca Corrente* Grants 2022-2024 and by *Paolo Zorzi Association for Neuroscience* (2024-2026 EPICARE project). We thank Y. Yanagawa (Gunma University, Japan) for providing GAD67 knock-in mice and G. Szabo (Inst. Experimental Medicine, Budapest, Hungary) for providing GAD65 transgenic mice.

## Author contributions

P.S. planned the experiments and was responsible for the interpretation of the findings; he performed the experiments and the data analysis and contributed to write the manuscript; C.M. and V.G. contributed to perform experiments and to data analysis; L.U. performed the experiments on the *in vitro* isolated guinea pig brain; M.C.R. performed the immunohisochemical experiments and provided technical support for the preparation of the experiments; M.d.C. contributed to experimental planning, data interpretation and manuscript writing; he also provided the funding to run the experimental activities.

The Authors declare no competing interests.

